# Hemispheric decoupling of awareness-related activity in human auditory cortex under informational masking and divided attention

**DOI:** 10.1101/2023.03.20.533547

**Authors:** Andrew R. Dykstra, Alexander Gutschalk

**Affiliations:** Department of Biomedical Engineering, University of Miami, Coral Gables, FL, USA; Department of Neurology, Ruprecht-Karls-Universität Heidelberg, Heidelberg, Germany

**Author notes:** Corresponding author, Andrew R. Dykstra, Department of Biomedical Engineering, University of Miami.

**Keywords:** attention, awareness, auditory cortex, multitone masking, perception

## Abstract

The conditions under which sensory stimuli require selective attention to reach awareness is a fundamental question of cognitive neuroscience. We examined this question in the context of audition utilizing M/EEG and a dual-task informational-masking paradigm. Listeners performed a demanding primary task in one ear – detecting isochronous target-tone streams embedded in random multi-tone backgrounds and counting within-stream deviants – and retrospectively reported their awareness of secondary, masker-embedded target streams in the other ear. Irrespective of attention or ear, left-AC activity strongly covaried with target-stream detection starting as early as 50 ms post-stimulus. In contrast, right-AC activity was unmodulated by detection until later, and then only weakly. Thus, under certain conditions, human ACs can functionally decouple, such that one – here, right – is automatic and stimulus-driven while the other – here, left – supports perceptual and/or task demands, including basic perceptual awareness of nonverbal sound sequences.

## INTRODUCTION

The conditions under which sensory stimuli must be attended in order to reach awareness is a fundamental question of cognitive science. Although selective attention and conscious perception have long been thought to be inextricably linked^1–3^, recent work has suggested that they may be dissociable^4^ [see also^5,6^]. However, as for conscious perception itself^7^, knowledge concerning the relationship between it and attention comes primarily from vision. It thus remains unclear how such knowledge might apply to audition, particularly given the different organism-level functional roles served by the two systems^8,9^, i.e., audition’s role as an “early-warning system”^10^.

Previous work examining neural correlates of auditory perceptual awareness identified the awareness-related negativity^9,11–14^ (ARN), alternatively termed the auditory awareness negativity^15^ (AAN). The AAN is a broad negativity between approximately 75 and 300 ms associated with perceptual awareness of target acoustic stimuli (tones, typically), and can be thought of as an auditory version of a perceptual awareness negativity^16^, with counterparts in both the visual and somatosensory domains. However, some have questioned whether the AAN (and analogues in other domains) might reflect correlates of selective attention^17–20^ rather than perceptual awareness. Indeed, the AAN overlaps with other components known to be strongly modulated by attention^21–27^ in both space and time [but see ^12^]. It thus remains unclear whether neural correlates of auditory perceptual awareness can be dissociated from those of selective attention, either for the AAN or other putative correlates of conscious perception in audition.

Here, we address these questions using a dichotic informational masking paradigm and simultaneous M/EEG. Listeners performed a demanding dual-task in one ear – detecting regularly repeating pure-tone target streams embedded in multi-tone masker clouds (Fig. 1), and subsequently counting amplitude-modulated deviants within the target stream – and retrospectively reported whether they also detected similar masker-embedded target streams in the other, secondary ear. For both primary- and secondary-ear targets, M/EEG responses were binned according to listeners’ perceptual report (i.e. detected vs. undetected targets, equalized for number of trials), averaged, and mapped to the cortical surface using noise-normalized minimum-norm estimates.

**Figure 1.**
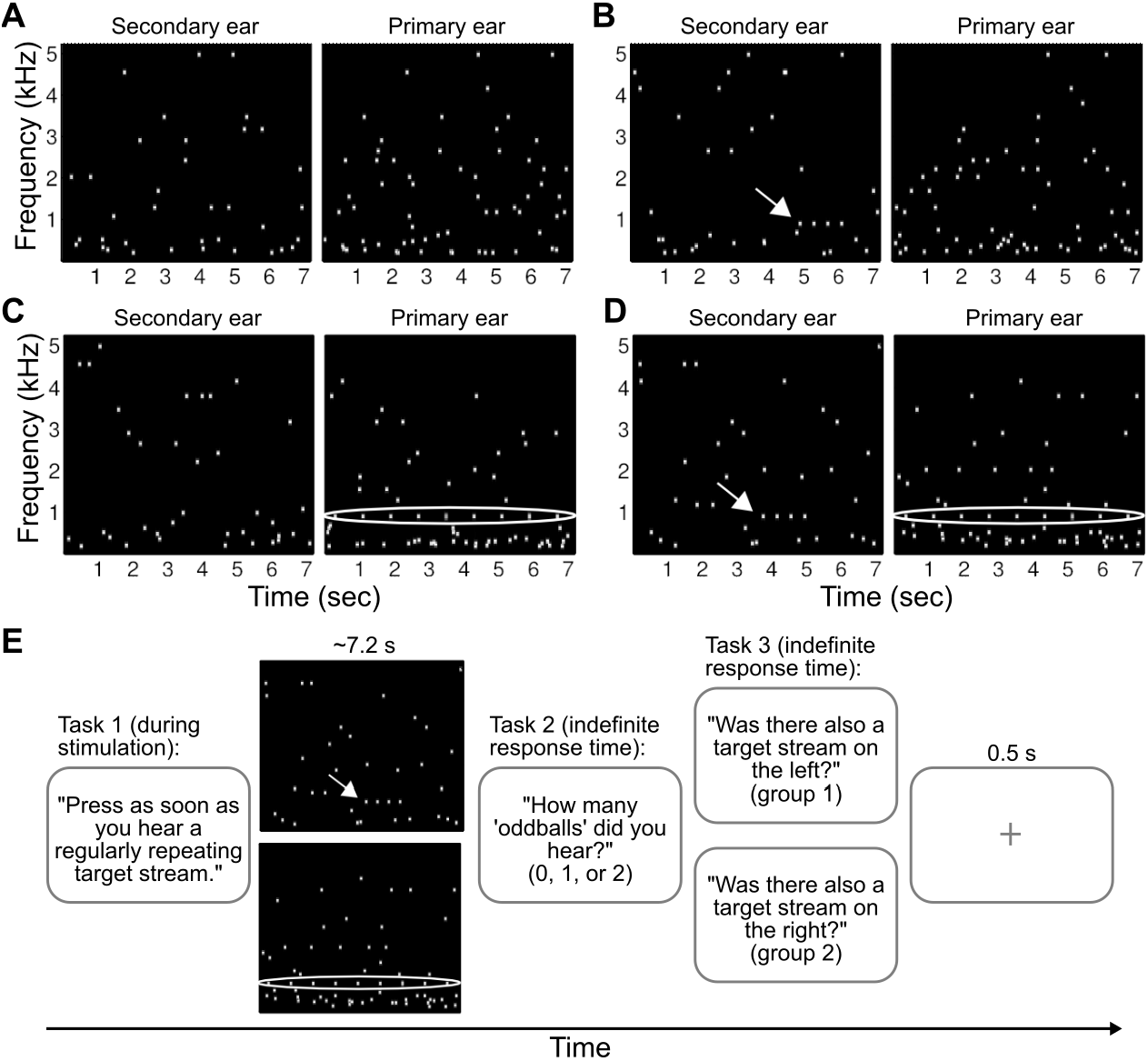
Stimuli and trial structure. **(A-D)** Spectrograms from example primary- and secondary-ear stimulus sequences. **(A)** No target stream present on either the primary or secondary ears. (B) Primary-ear target stream not present, secondary-ear target stream present. **(C)** Primary-ear target stream present, secondary-ear target stream not present. **(D)** Both primary- and secondary-ear target streams present. (E) Trial structure. On each trial, listeners had three tasks: (i) to listen to the designated (that is, primary) ear and press a button as soon as they began to hear a regularly repeating pure-tone target stream, (ii) to count the occurrence of amplitude-modulated deviant tones within the target stream, and (iii) to report whether they also happened to hear a brief, isochronous target stream on the other (that is, secondary) ear.

Because the study was initially focused on the relationship between attention and awareness in audition and strong asymmetry of M/EEG responses was not expected, the primary and secondary ears were initially not counterbalanced. That is, in the first group of listeners, the primary (secondary) ear was always the right (left). However, inspection of this group’s M/EEG data revealed a strongly left-dominant pattern of assumed awareness-related activity in auditory cortex. We therefore repeated the experiment in a second group of listeners with the ears flipped, i.e., the primary (secondary) ear was now the left (right). This allowed us to examine whether the lateralization effects we observed in group 1 were reflective of a true hemispheric asymmetry or rather a product of the focus of auditory spatial attention, as has been observed for certain lateralized auditory evoked responses previously^22,23^. If the latter were true, we’d expect the lateralization pattern to be reversed between groups 1 and 2, whereas the former interpretation would predict that it would remain the same.

## METHODS

All procedures were approved by the Institutional Review Board at the University of Heidelberg, and participants gave written informed consent prior to participation.

### Listeners

32 healthy participants between the ages of 21 and 30 (10 female) were recruited for the study. None of the listeners reported any history of hearing disorders. Six additional subjects were excluded from analysis due to poor behavioral performance (N=4) or excessive artifact in the MEG (N=2).

### Stimuli and Procedure

All stimuli were generated in MATLAB (The Mathworks, Natick, MA) and stored as 32-bit wave files at a sampling rate of 48 kHz. The wave files were converted to analog waveforms by an on-board sound card (RME DIGI96/8 PAD) and freestanding DA converter (RME ADI-8 DS, RME, Haimhausen, Germany), which in turn were controlled by SoundMexPro software (SoundMexPro, Oldenburg, Germany) in the MATLAB environment. The analog stimuli then passed through a programmable attenuator (TDT PA5) before being amplified (TDT HB7, Tucker-Davis Technologies, Alachua, FL, USA) and presented to listeners via ER-3 insert earphones (Etymotic Research, Elk Grove Village, IL, USA).

The stimuli used for the study were sequences of short-duration pure tones placed randomly in time and frequency^11–13^ (Fig. 1, supplemental audio files). Each sequence was approximately 7.5 seconds in duration and determined independently for each ear. Individual tones were 100 ms in duration (with 10 ms on- and off-ramps) and drawn from a log-uniform distribution of frequencies between 200 and 5000 Hz. In each ear, sequences were comprised of either masker tones alone or masker tones plus a regularly-repeating (isochronous) target-tone stream. Sequences in the primary ear were defined in successive 800-ms windows in which seven masker tones with onsets occurring at any point within the window were presented. On target+masker trials, one of the masker tones in each window was replaced by the target such that the within target-stream onset asynchrony (SOA) was 800 ms. For each trial with a target, target tones had a frequency of either 585, 914, or 1430 Hz, and were surrounded by a spectral protected region which prevented the occurrence of masker tones in the range 0.8*F_targ_ – 1.25*F_targ_. where F_targ_ is the frequency of the target tones.

Stimuli in the other, secondary ear were defined in similar fashion, but in successive 400-ms (rather than 800-ms) windows in which two masker tones (vs. seven for the primary ear) with onsets occurring at any point within the window were presented. When a target stream was present, it consisted of four identical tones with an SOA of 400 ms. As for the primary ear, secondary-ear target frequencies were either 585, 914, or 1430 Hz (with equal likelihood across all classes of primary-ear trials), and were surrounded by the same protected region. We used such brief targets with a sparse masker cloud in the secondary ear for two reasons, (i) to increase the salience of the secondary-ear targets and (ii) to make the retrospective task one of reporting a unitary event (the presence of the four tones constituting the target stream) instead of an ongoing task as for the primary ear.

Because we did not expect strongly asymmetric results and because this was not the initial focus of the study, in a first group of listeners (N=16), the primary and secondary ears were not counterbalanced, i.e., the right (left) ear was always primary (secondary). However, the responses turned out to be very asymmetric (cf. Figs. 3–6), and we subsequently repeated the experiment in a second group of listeners (N=16) with the primary and secondary ears switched in order to test whether the lateralization present in the first group was reflective of a true hemispheric asymmetry or rather a product of the focus of auditory spatial attention. If the latter were true, we’d expect the lateralization pattern to be reversed between groups 1 and 2, while the former interpretation would predict it to remain the same.

### Data Acquisition

MEG signals were recorded with an Elekta Neuromag 122-channel whole-head system comprised of two, orthogonal planar gradiometers at each of 61 locations around the head^28^ inside a 4-layer electromagenetically-shielded room (IMEDCO AG, Hägendorf, Switzerland). EEG was recorded simultaneously via a 63-channel mesh cap (EasyCap GmbH, Herrsching, Germany) with equidistant channel spacing. Both MEG and EEG were acquired via the Neuromag recording system at a sampling rate of 500 Hz and an online low-pass filter with a cutoff frequency of 160 Hz. Stimulus trigger pulses were produced by the remaining 6 channels of the same 8-channel sound card used to present the sound stimuli to the participant and also recorded by the Neuromag acquisition system. Thus, temporal resolution of stimulus-locked averaging was limited only by the 500-Hz sampling rate of the M/EEG acquisition. Participants gave their behavioral responses with their dominant hand (15 right, 1 left) via an optical button device (Current Designs, Philadelphia, PA, USA) whose key strokes were read via USB and aligned with the M/EEG recording offline.

Prior to each M/EEG session, the locations of each EEG electrode as well as 15 additional head points, referenced to the coordinate system defined by three cardinal points (left and right periauricular points, and the nasion), were digitized using the Polhemus tracker system (Colchester, Vermont, USA). Participants’ head position within the dewar was measured by four, head position-indicator coils placed on the left and right mastoid and forehead prior to commencing the experiment, and participants were instructed to remain as still as possible during the approximately 50 minute-long recording session (split into two equally long blocks).

In order to create an individualized source space and boundary-elemant model for each participant, we additionally acquired three anatomical MRI sequences via a 3-Tesla TIM Trio scanner from Siemens (Erlangen, Germany). The first was a high-resolution (1 mm isotropic), T1-weighted scan (MPRAGE), and the other two were T1-weighted, multi-echo FLASH sequences with 5- and 30-degree flip angles and the same field of view and resolution as the MPRAGE acquisition.

### Data Preprocessing

Data from each block were inspected for time epochs and/or channels containing large artifacts or flat signal (such data were rejected). Because no EOG was recorded, we synthesized such a signal from three frontal EEG channels immediately above the eye sockets, and used this signal in order to define and add blink event markers to the raw files. In order to mitigate the contribution of ocular and cardiac signals from affecting the time-locked averages, independent component analysis, as implemented by the binica algorithm in EEGLAB^29^, was run on each block of raw data (separately for MEG and EEG). Components comprised predominantly of blinks, saccades, and cardiac pulsation were identified by visual inspection and removed from the ICA weights matrix before back-projection.

The raw data were then passed through zero-phase shift IIR filters to suppress activity outside the 0.5-15 Hz passband, divided into epochs spanning −100-700 ms (primary-ear targets) or −100-1700 ms (secondary-ear targets) relative to target-tone or target-stream onset for primary- and secondary-ear targets, respectively. After ICA removal of blinks, saccades, and cardiac activity, any epoch still containing EEG signals greater than ±150 μV or gradiometer signals greater than ±10 pT/cm was rejected. The epochs were then binned according to whether the target tones were detected or not. Additional epochs, comprised of “virtual targets,” were created from the catch-trial (i.e., masker-alone) stimuli by time-locking the M/EEG signal relative to virtual target tones, i.e. the positions in the sequence of potential target tones, had they been present. This condition serves as a baseline with which to compare the responses to both detected and undetected targets and, because the time-locking used to create the averaged response is essentially random, the resulting average should be flat, thus yielding an empirical estimate of the averaging procedure’s signal-to-noise ratio. Prior to averaging and baseline correction, event counts across conditions were equalized such that the temporal distribution of epochs is (approximately) the same across conditions. This procedure is necessary prior to computing noise-normalized source estimates to avoid biased z-scores and to mitigate possible time-varying noise characteristics during the recording.

Finally, we created masker-tone epochs via a time-frequency based procedure as described previously^12^. Briefly, masker-tone events were obtained by computing spectrograms for each channel of every stimulus sequence and subsequently identifying energy onsets in each frequency band (after filtering out the band containing the target-tone stream, when present). Parameters – 768-sample Hanning windows, 87.5% overlap, 16,384-point FFTs – were chosen such that the temporal resolution of the resulting spectrograms was 2 ms, ensuring event timing that was as least as resolved as the 500-Hz M/EEG acquisition. The resulting masker-tone events were sorted according to (i) whether they came from the primary or secondary ear, (ii) target-tone detection (detected vs. undetected trails), and (iii) early vs. late portions of the trial. For targets-detected trials, the early and late periods corresponded to before and after detection, respectively. For targets-undetected trials, masker events were deemed early (late) if they occurred before (after) the average detection time from targets-detected trials. For primary-ear maskers, epochs spanned from 100 ms before to 700 ms after masker-tone onset. For secondary-ear maskers, this range spanned from −50 to 400 ms.

High-resolution T1-weighted MRIs were used to create 3D reconstructions of the white and pial surface of each participant using Freesurfer^30–32^ (http://surfer.nmr.mgh.harvard.edu/). Using the MNE suite, an anatomically-constrained M/EEG source space was created by decimating the resulting white matter surface to ~4100 dipole locations per hemisphere, resulting in each location representing an approximate surface area of 24 mm2, with inter-source spacing of ~5mm. The multi-echo FLASH sequences were used to produce high-resolution inner- and outer-skull and scalp surfaces. These, in turn, were decimated and used to create realistic 3-layer boundary-element head models (BEM), which has been shown to result in a more accurate forward model (i.e. gain matrix), and thus better localization of EEG sources with little, if any, impact on the accuracy of forward modeling and source reconstruction of MEG data^33,34^. Finally, the MEG and MRI coordinate frames were aligned by first manually demarking the fiducial points utilizing the high-resolution scalp surface. Refinement of this initial 3D affine alignment was carried out using the iterative closest-point algorithm^35^ as implemented by the MNE suite.

### Source Estimation

We utilized anatomically-constrained, noise-normalized minimum-norm estimates^36,37^, also known as dynamic statistical parametric maps, or dSPMs^38^. For each participant (N=32), a dSPM estimate of the averaged M/EEG response to detected targets was computed, with fixed dipole orientations (i.e. dipoles normal to the white-matter surface) and no depth weighting. Noise covariance matrices were estimated from the concatenation of prestimulus baseline epochs (from 100 – 2 ms preceding stimulus onset) across all conditions. The dSPMs from each participant were then mapped to the Freesurfer average brain using a 2D spherical registration method known to yield accurate alignment of functional and anatomical brain areas across individuals^31^. This resulted in the grand-average source maps shown in Figs. 3 and 5; note that while these maps are the result of a fixed-effects analysis (i.e. simply averaging across individual listener’s responses), all significance tests were performed on the source waveforms and amplitude quantifications used random-effects models. These maps indicated that the bulk of activity observed in the sensor-level averages likely arose from auditory cortex on the superior temporal plane. Grand-average source waveforms were computed by averaging individual source-level activity across vertices within a region of interest defined by the transverse temporal gyrus and planum temporale (i.e., the posterior aspect of the superior temporal plane, or pSTP).

### Statistical Analysis

Behavioral data as well as component peak amplitudes/latencies were assessed through the use of repeated-measures analysis of variance (ANOVA) and paired t-tests. dSPM source waveforms from detected and undetected targets were contrasted using a modified version of the non-parametric cluster-based tests outlined by Maris and Oostenveld^39^. This method has proven to be an effective way to mitigate the multiple comparisons problem, and allows for delineation of the extent of significant differences between conditions. Because we conducted such tests on individual waveforms (rather than across a surface or in a volume), it was necessary only to control for multiple comparisons in the temporal dimension. Because we hypothesized differences between detected targets and undetected targets, we assigned a cluster-wise alpha level of 0.05, and took any differences with p-values less than this threshold to be statistically significant.

## RESULTS

To examine neural correlates of consciously perceived sounds outside the focus of selective attention, we recorded combined M/EEG while listeners (N=32) engaged in a dichotic, multi-task informational-masking paradigm. Spectrally isolated suprathreshold target-tone streams were sometimes rendered inaudible by embedding them in random multi-tone masker clouds (Fig. 1, supplemental audio files). On each trial, listeners were instructed to attend to a designated ear and indicated via button press the moment at which they began to hear out the isochronous target stream from the multi-tone background. Upon detecting the target stream, listeners were then required to count the occurrence of amplitude-modulated deviants within it, thus ensuring that their attention was fixed upon the designated side for the duration of each trial. Finally, upon the cessation of each sequence, listeners were also asked to report whether they had incidentally heard a similar masker-embedded target stream in the other, secondary ear.

### Behavior

For the primary ear, detection rates for the masker-embedded target streams plateaued at 0.75 ± 0.03 (group 1) and 0.77 ± 0.02 (group 2) (Fig. 2A). False-alarm rates, while also increasing throughout the stimulus sequence, remained low overall, never exceeded 0.10 ± 0.02 (group 1) and 0.06 ± 0.01 (group 2), resulting in sensitivity (d’) that plateaued at 2.1 ± 0.1 (group 1) and 2.4 ± 0.1 (group 2) (Fig. 2B). Similarly, d’ was always significantly higher than zero (group 1: *T*_15_ ≥ 4.0, *p* ≤ 0.001, group 2: *T*_15_ ≥ 6.9, *p* ≤ 0.001), and increased with increasing tone position (group 1: *F*_8,120_ = 96.2, *p* ≪ 0.001, group 2: *F*_8,120_ = 54.6, *p* ≪ 0.001), indicating that listeners were sensitive to the presence of the target streams, particularly for later tone positions. Group 2 performed slightly better overall (*F*_1,30_ = 8.7, *p* < 0.01).

**Figure 2.**
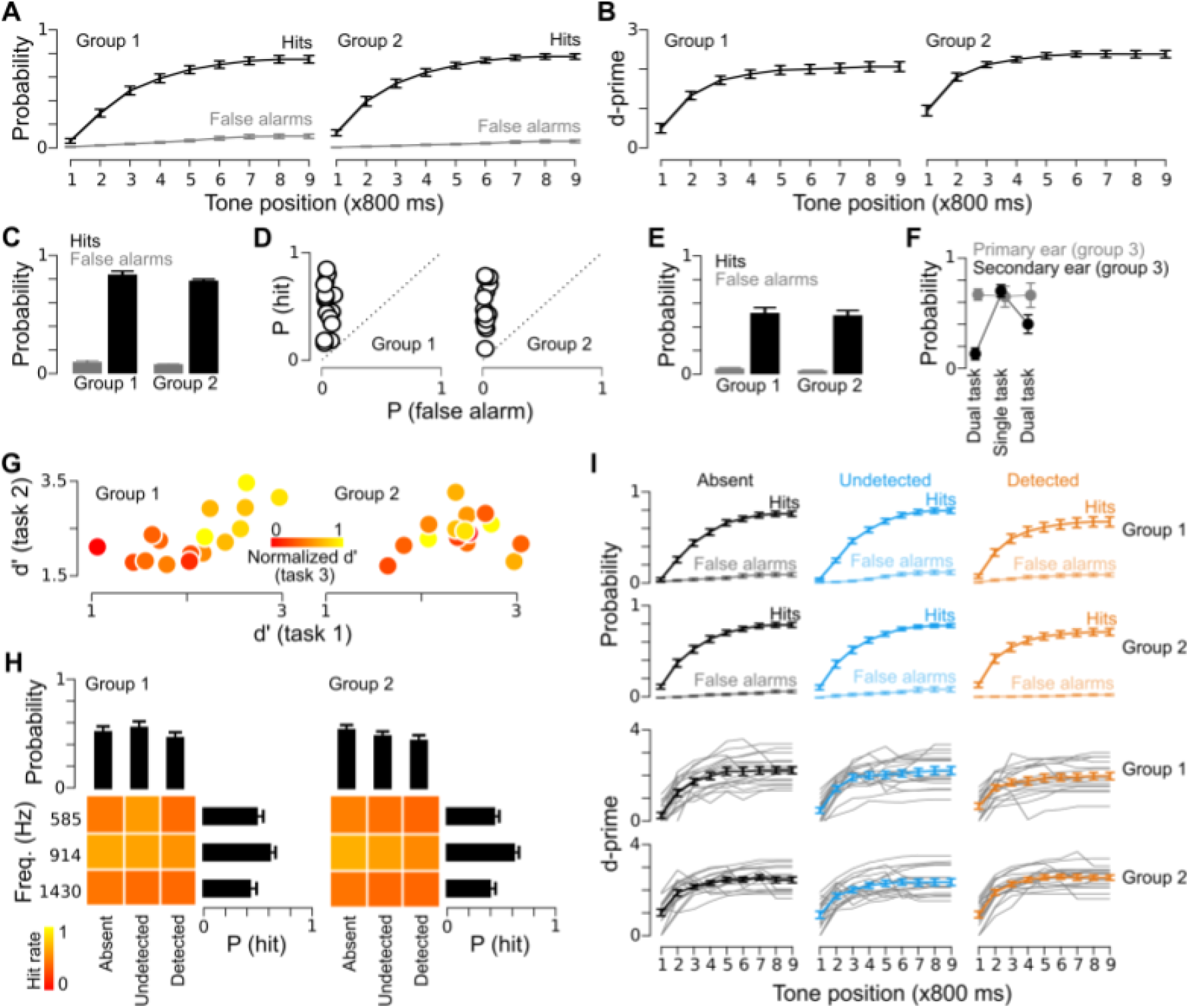
Behavioral results. **(A)** True- and false-positive rates for the primary-ear task (detect isochronous target streams embedded in multitone maskers) as a function of position within the stimulus sequence. (B) d’(z-scored true-minus false-positive rates) as a function of position within the stimulus sequence. (C) True- and false-positive rates for the second primary-ear task (count the occurrence of amplitude-modulated deviants within the isochronous target stream). (D) True-vs. false-positive rates for the secondary-ear task. (E) Same data as in D, averaged across listeners. (F) Behavioral performance of group 3. (G) Relationship between performance on primary- and secondary-ear tasks. **(H)** Secondary-ear performance sorted by primary-ear percepts. **(I)** Primary-car performance sorted by secondary-car percepts.

**Figure 3.**
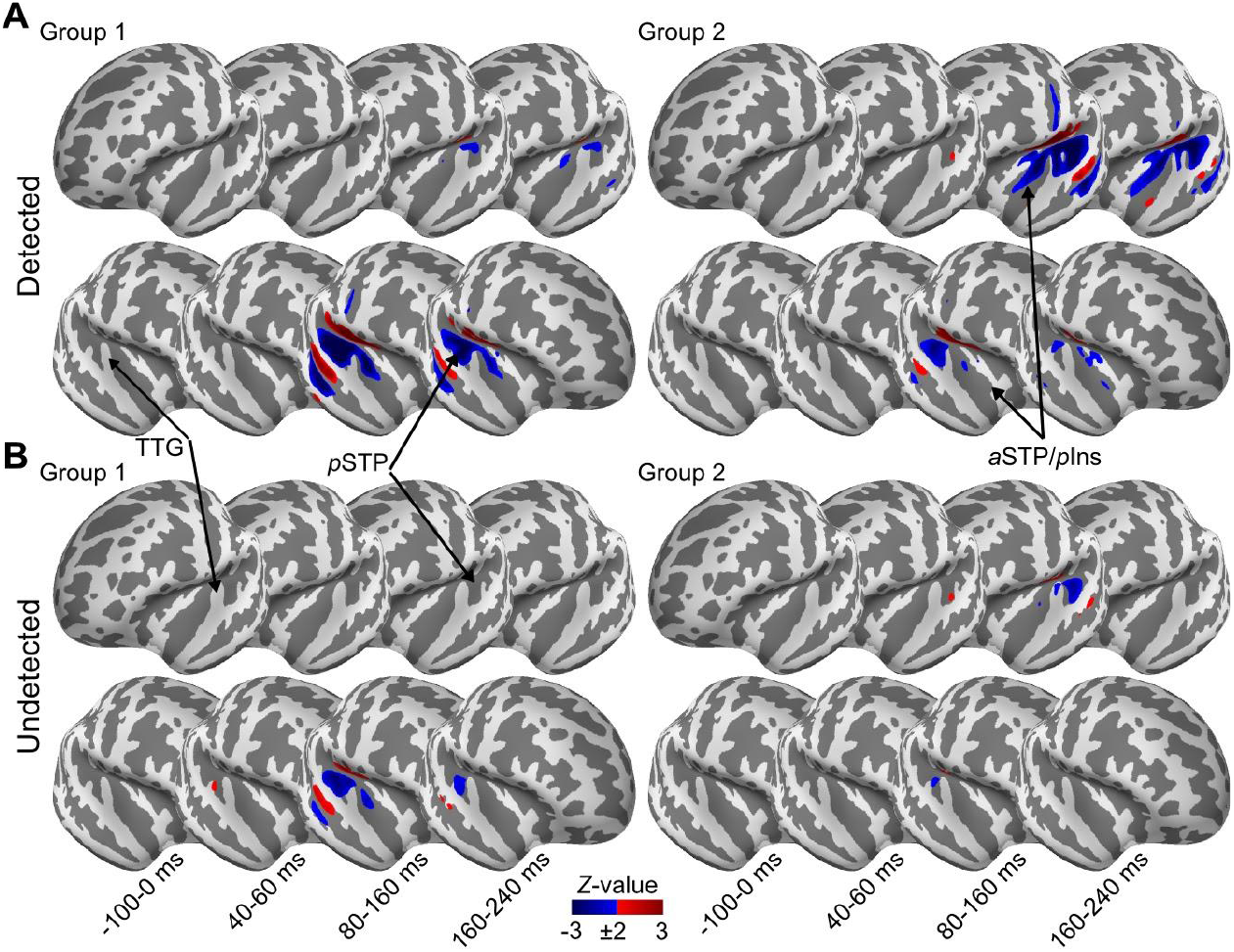
Whole-brain activity maps for secondary-ear targets. **(A)** Grand-averaged distributed source estimates for detected, unattended-side targets in each of four time windows (−100-0 ms, 40-60 ms, 80-160 ms, and 160-240 ms). (B) Same as in (A), for undetected targets. TTG: transverse temporal gyrus. *p*STP: posterior superior temporal plane. *a*STP: anterior superior temporal plane. *p*Ins: posterior insula.

**Figure 4.**
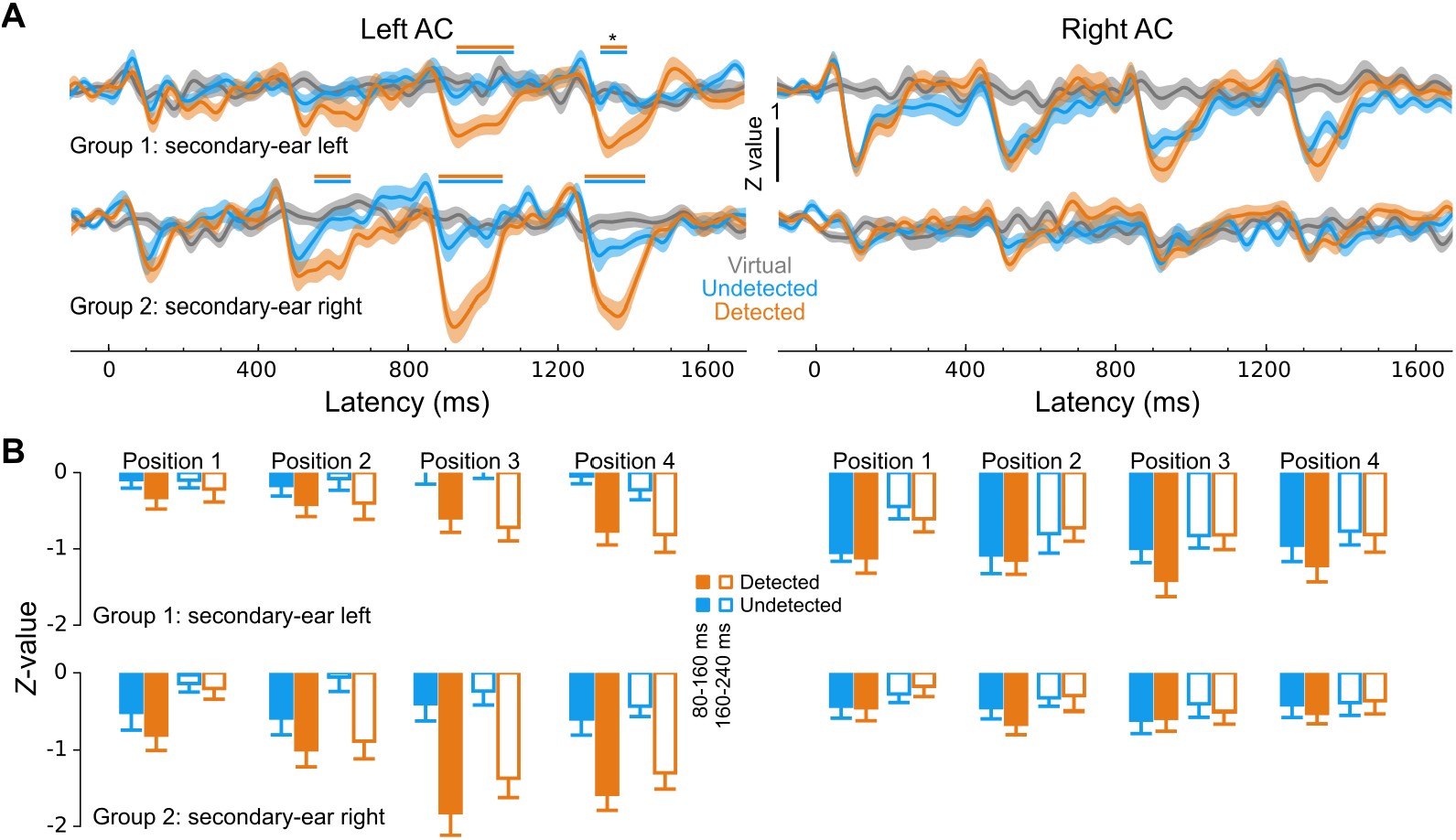
Secondary-ear time courses and peak amplitudes from posterior auditory cortex. **(A)** Grand-averaged timecourses for each group separately for detected (orange), undetected (blue), and virtual (gray) targets. Horizontal bars above the traces denote cluster-level significantly different (*p* < 0.05) responses between detected- and undetected-target responses (*: *p* = 0.06). **(B)** Peak amplitudes for detected (orange) and undetected (blue) targets for both early (80-160 ms) and late (160-240 ms) time windows (closed and open bars, respectively) for each target-tone position.

**Figure 5.**
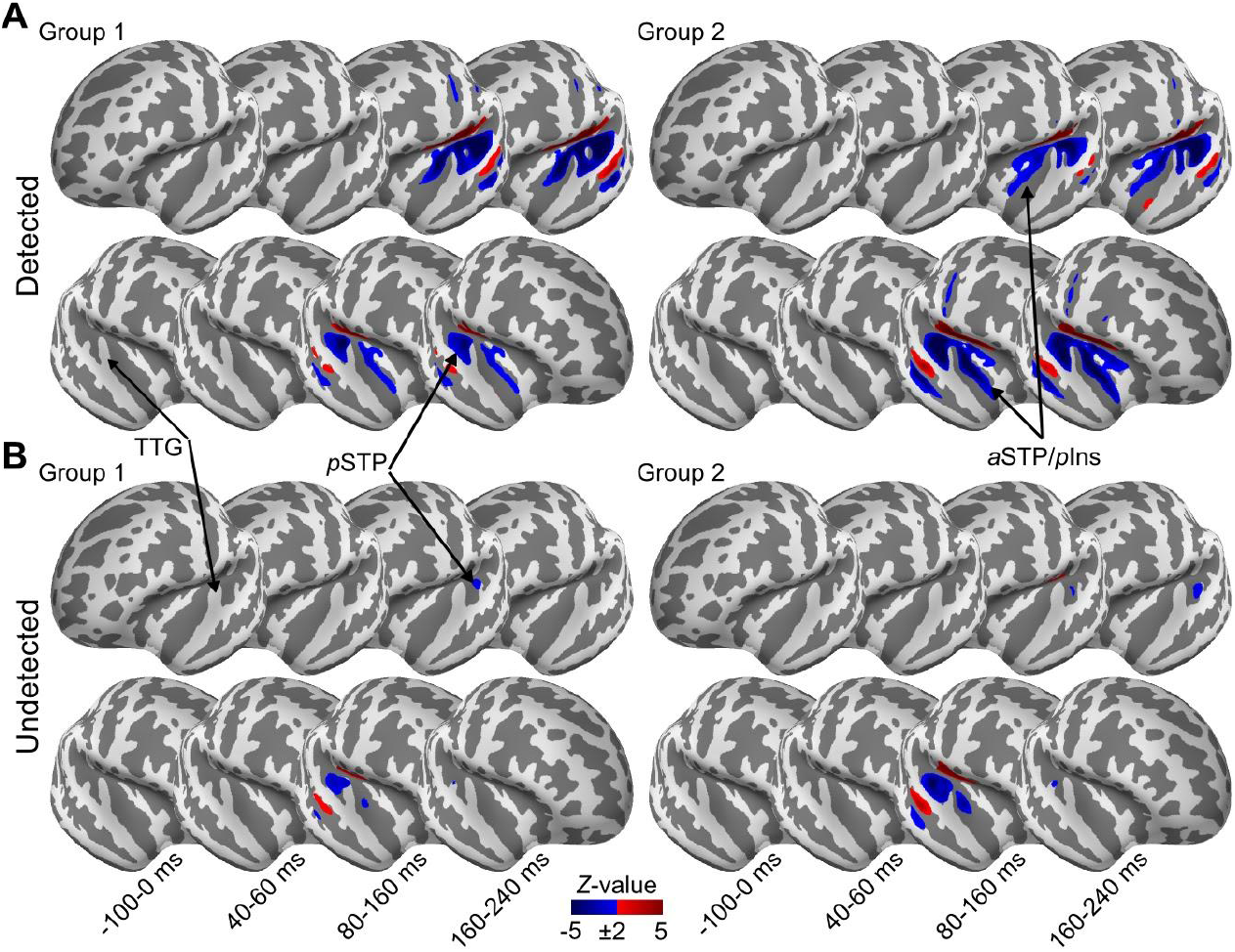
Whole-brain activity maps for primary-ear targets. **A)** Grand-averaged distributed source estimates for detected, primary-ear targets in each of four time windows (−100-0 ms, 40-60 ms, 80-160 ms, and 160-240 ms). (B) Same as in (A), for undetected targets. Abbreviations as in Fig. 3.

**Figure 6.**
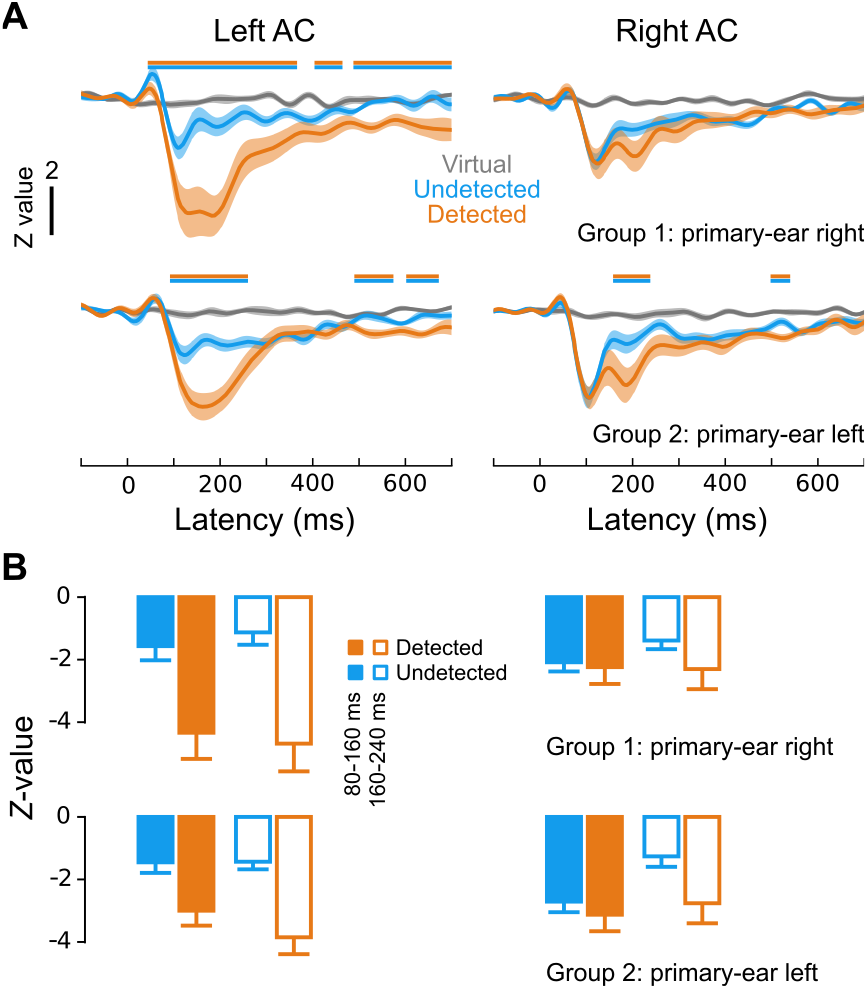
Primary-ear time courses and peak amplitudes from posterior auditory cortex. **(A)** Grand-averaged timecourses for each group separately for detected (orange), undetected (blue), and virtual (gray) targets. Horizontal bars above the traces denote cluster-level significantly different (*p* < 0.05) responses between detected- and undetected-target responses. (B) Peak amplitudes for detected (orange) and undetected (blue) targets for both early (80-160 ms) and late (160-240 ms) time windows (closed and open bars, respectively).

To ensure that listeners maintained attention toward the designated side, their second primary-ear task was to count the occurrence of amplitude-modulated deviants within detected target streams (either one or two) (Fig. 2C). For both groups, hit rates (group 1: 0.83 ± 0.04, group 2: 0.78 ± 0.02) were substantially higher than false-alarm rates (group 1: 0.09 ± 0.02, group 2: 0.07 ± 0.01) (group 1: *T*_15_ =21.2, *p* ≪ 0.001, group 2: *T*_15_ = 36.4,*p* ≪ 0.001). These values resulted in d’ values of 2.4 ± 0.1 (group 1) and 2.2 ± 0.1 (group 2), which did not differ between groups (unpaired t-test: *T*_30_ = 1.3, n.s.). Importantly, hit rates for the deviant-detection task did not reach ceiling, suggesting that listeners’ attention was strongly engaged by the primary-ear tasks. Querying listeners after the experimental session also gave this impression.

In spite of this, they were also sensitive to the presence of brief, sparsely masked target streams presented to the secondary ear (Fig. 2D & 2E). On average, hit rates were 0.51 ± 0.05 (group 1) and 0.49 ± 0.05 (group 2), with substantial variability across participants typical of informational masking paradigms^40^. False-alarm rates remained low (group 1: 0.04 ± 0.01, group 2: 0.03 ± 0.01), never exceeding 0.13 (group 1) or 0.07 (group 2), resulting in d’ values of 1.8 ± 0.2 (group 1) and 1.9 ± 0.1 (group 2), which were significantly higher than zero for both groups (group 1: *T*_15_ = 9.9,*p* ≪ 0.001, group 2: *T*_15_ = 13.4,*p* ≪ 0.001), and did not differ between groups ( *T*_15_ = 0.35, *n.s*.). Importantly, despite the fact that the secondary-ear masker clouds were much sparser than their primary-ear counterparts (cf. Fig. 1, Methods), d’ was smaller for secondary-ear targets, again suggesting that listeners attended the instructed side.

To examine explicitly whether the primary-ear task was engaging enough to affect performance on the secondary-ear task, we asked 10 additional listeners (group 3) to perform a behavioral experiment in which sensitivity to nominally secondary-ear targets was examined in the absence of the primary-ear task (Fig. 2F). In order, each listener in this third experiment performed (i) the same dual task as in groups 1 and 2, (ii) the “primary-ear” task alone (while ignoring “secondary-ear” stimuli), (iii) the “secondary-ear” task alone (while ignoring “primary-ear” stimuli), and (iv) the dual task. A 2-way repeated-measures ANOVA with ear (primary vs. secondary) and block (dual task vs. single task vs. dual task) as factors revealed a significant ear by block interaction (*F*_2,18_ = 54.48, *p* < 0.001). While performance on the “primary-ear” task did not change with condition (*F*_2,18_ = 0.083, *n.s*.), performance on the secondary-ear task dramatically improved when listeners were not simultaneously performing the primary-ear task (*F*_2,18_ = 74.83, *p* < 0.001). This pattern of behavioral results for this group indicates that our attention manipulation was effective, such that “secondary-ear” targets were truly secondary, and less attended than their “primary-ear” counterparts.

To further examine whether listeners maintained their attentional focus as instructed, we examined the relationship between primary- and secondary-ear behavioral performance (Fig. 2G). If listeners let their attention drift towards the secondary ear, we would expect an inverse relationship between primary- and secondary-ear performance. However, this is not what we observed. In fact, group 1 displayed the opposite pattern, i.e. positive correlations between primary- and secondary-ear performance (Task 2 vs. Task1: *R*_15_ = 0.70, *p* < 0.01; Task 3 vs. Task 1: *R*_15_ = 0.83, *p* < 0.001; Task 3 vs. Task 2: *R*_15_ = 0.70, *p* < 0.01), indicative of a participant-specific effect. Group 2 showed no significant correlations between any of the three tasks. Critically, correlations between performance on the secondary-ear task (task 3) and either of the primary-ear tasks (tasks 1 and 2) were never negative.

To further examine the relationship between primary- and secondary-ear performance, we sorted the secondary-ear data according to primary-ear percept (i.e., whether the primary-ear targets were detected, undetected, or absent) (Fig. 2H), and vice versa (Fig. 2I). If listeners let their attention drift towards the secondary ear, we’d expect secondary-ear performance to come at the expense of primary-ear performance, and vice versa, but this is not what we observed. For the secondary ear (Fig. 2H), hit rates were significantly affected by primary-ear percepts (generally higher when the primary-ear targets were either subliminal or absent) (*F*_2,60_ = 5.2, *p* < 0.01). However, this effect was small, much less robust than the effect of target-stream frequency (*F*_2,60_ = 18.2, *p* ≪ 0.001), and not present for all target-stream frequencies (interaction between primary-ear percept and secondary-ear frequency: *F*_4,120_ = 2.5, *p* < 0.05; also see Table S1).

For the primary ear (Fig. 2I), neither hit rates nor d’ were significantly affected by secondary-ear percept (hit rates: *F*_2,60_ = 2.1, *n.s*., d’: *F*_2,60_ = 0.13, *n.s.*), though there was a highly significant interaction between secondary-ear percept and tone position for hit rates (*F*_16,480_ = 25.6, *p* ≪ 0.001), indicating that secondary-ear percepts may have reduced primary-ear hit rates, at least for later tone positions (Table S2). Again, however, this effect was small, only reached statistical significance when collapsing across the two groups, and did not translate to significantly different d’ values (*F*_16,480_ = 1.4, *n.s.*) for any of the nine tone positions (*F*_2,30_ ≤ 0.54, *p* ≥ 0.57), perhaps because a similar interaction between secondary-ear percept and tone position was also observed for false alarms (*F*_16,480_ = 4.1, *p* ≪ 0.001). However, subsequent one-way ANOVAs on false-alarm rates at each of the nine tone positions did not reach statistical significance (*F*_2,30_ ≤ 2.1, *p* ≥ 0.13). In summary, detection of secondary-ear targets did seem to briefly distract listeners from their primary-ear tasks, suggesting that the secondary-ear targets captured attention momentarily, though the effects were small.

### M/EEG: Secondary Ear

Neural responses to secondary-ear target streams were analyzed using anatomically constrained, noise-normalized minimum-norm estimates^36–38^ (Figs 3–4). Due to the fact that listeners were to report the presence/absence of the secondary-ear target streams at the end of each trial, we strove to make them brief, unitary events, such that listeners were able to make a simple “yes or no” response. We thus analyzed the response to secondary-ear target streams across all four tones in the sequence, with the exception of the grand-averaged statistical maps of Fig. 3, where all tones were collapsed into a single response to examine their source profiles. The results indicate that left-AC activity robustly differentiated between detected and undetected targets in both groups, though this effect was stronger in group 2, for whom the secondary-ear targets were presented to the right ear. In contrast, while right-AC activity was much larger in response to left-vs. right-ear targets, it did not differentiate between detected and undetected targets in either group.

We quantified these effects by computing source-amplitude values for each group, condition, hemisphere, time window (80-160 ms vs. 160-240 ms), and target-tone position (1 vs. 2 vs. 3 vs. 4) (Fig. 4). Four-way ANOVAs revealed four-way interactions between percept, hemisphere, time window, and tone position in each group, although this effect was only marginally significant in group 1 (group 1: *F*_3,45_ = 2.3, *p* = 0.09, group 2: *F*_3,45_ = 4.5, *p* < 0.01; cf. Table S3 for the full ANOVA results). Subsequent tests between responses at each combination of hemisphere, group, target-tone position, and time window indicated that (i) the responses elicited by detected and undetected targets never differed significantly in right AC, (ii) the earliest awareness-based difference emerged in left AC in the early time window (80-160 ms) at the first target-tone position, (iii) the left-AC difference between detected- and undetected-target responses increased in magnitude as a function of target-tone position, and (iv) responses in each group were overall larger contralateral to the ear of stimulation.

### M/EEG: Primary Ear

To examine whether this pattern of asymmetry was related to focus of attention, we then analyzed the responses to primary-ear targets (Figs. 5–6). As for secondary-ear targets, the primary source of activity elicited by primary-ear targets (both detected and undetected) was in auditory cortex (Fig. 5). Furthermore, in both groups, the primary-ear responses confirmed the strong pattern of left-dominant asymmetry observed for secondary-ear responses despite the fact that the stimuli generating the responses arose from different ears across the two groups. The source-level traces (Fig. 6A) confirm the general pattern shown in the maps, i.e., that at early latencies (beginning at 50 ms in group 1 and 96 ms in group 2), left-AC activity was much more strongly associated with perceptual awareness than was right-AC activity. Right-AC activity did show a difference between detected and undetected targets at longer latencies in both groups, but this effect did not reach statistical significance for group 1. Otherwise, right-AC activity, especially around the N1 time window (~80-160 ms), was remarkably similar – and large – in response to both detected and undetected targets.

To quantify these effects, we computed average source-amplitude values for each condition, hemisphere, and time window (80-160 ms vs. 160-240 ms) (Fig. 6B). The same general pattern holds (robust differentiation between detected and undetected targets in left AC, no differentiation in right AC at early latencies, weak differentiation in right AC at longer latencies). Three-way ANOVAs with percept (detected vs. undetected), hemisphere (right vs. left), and time window (80-160 ms vs. 160-240 ms) as factors revealed significant main effects of percept in both groups (group 1: *F*_1,15_ = 20.4, *p* < 0.001, group 2: *F*_1,15_ = 12.7, *p* < 0.01) as well as two-way interactions between (i) percept and hemisphere (group 1: *F*_1,15_ = 11.6, *p* < 0.01, group 2: *F*_1,15_ = 3.8, *p* = 0.07) and (ii) percept and time window (group 1: *F*_1,15_ = 8.0, *p* < 0.05, group 2: *F*_1,15_ = 33.8,*p* < 1e-4). We did not observe significant three-way interactions between percept, hemisphere, and time window for either group, as might be expected by visual inspection of Fig. 6 (i.e., a significant difference between detected and undetected targets in left AC for both time windows but only for the later time window in right AC; cf. Table S4 for the full ANOVA results). This is likely due to the fact that for both groups and both hemispheres, the difference between responses to detected and undetected targets was larger for the later time window, i.e., two-way interactions between percept and time window [Group 1: *F*_1,15_ = 3.6, *p* < 0.1 (left AC), *F*_1,15_ = 9.1, *p* < 0.01 (right AC); Group 2: *F*_1,15_ = 9.8, *p* < 0.01 (left AC), *F*_1,15_ = 18.7, *p* < 0.001 (right AC)]. However, the effect was more robust in group 2, and post-hoc paired t-tests confirmed that the detected/undetected difference was not significant in right AC for the earlier time window for either group, and only significant in group 2 for the later time window.

## DISCUSSION

Using a dichotic informational-masking paradigm and M/EEG, the present study demonstrated (i) that even sounds outside the focus of top-down attention have the capacity to reach awareness, (ii) that the neural correlates of such sounds are similar to those of their more attended counterparts, and (iii) a robust hemispheric asymmetry of awareness-related activity in human auditory cortex under certain conditions. The findings have several important implications for neurally based models of perceptual awareness, selective attention, and hemispheric asymmetry.

### Sounds Initially Outside the Focus of Selective Attention Can Reach Awareness

To the extent that our secondary-ear target sequences were outside the focus of selective attention, our findings argue against the notion of a hard top-down attentional requirement for conscious perception, consistent with certain previous results from vision (for review see ^41^). Certainly, our “secondary-ear” targets were not as well attended as their “primary-ear” counterparts (cf. Fig 2). They also suggest that the AAN, or at least some portion of it, is associated with perceptual awareness rather than reflecting response enhancement that is primarily related to the focus of selective attention^12^(also see Figs S3 and S4) despite its spatiotemporal overlap with attention-related components^21–27^ (but see ^20^).

However, we cannot rule out the possibility that the secondary-ear sequences, which were task relevant, “captured” listeners’ attention in a bottom-up, saliency-driven manner^42^ and subsequently triggered an attentional reorienting, such that later tones in the secondary-ear sequences were momentarily within the focus of selective attention. This would still be consistent with the interpretation of the perceptual awareness negativity reflecting selective attention rather than awareness^20^. In addition, the fact that secondary-ear targets were task relevant may have also played a role in biasing listeners’ strategy, perhaps putting them in a “split” attentional state^43,44^, directing residual attentional resources toward the secondary ear. However, the spotlight of auditory attention has long been thought to be contiguous^45,46^ (but see ^47^), and removing the secondary-ear task did not significantly improve performance on the primary task (Fig. 2F).

What is certain is that listeners’ attention was strongly engaged by their primary-ear tasks at the moment the secondary-ear sequences were presented. This is reflected by the fact that despite a relatively sparse secondary-ear masker cloud, only ~50% of secondary-ear targets were perceived compared to nearly 80% detection of primary-ear targets (or 60% after a similar absolute number of target tones). It is further reflected by the fact that listeners were much more sensitive to the presence of nominally secondary-ear targets when no primary-ear task was performed.

Thus, target sounds that are not strongly within the focus of selective attention (at least not a priori) can still reach awareness. In a previous study with a dichotic distraction of attention, participants did not retrospectively report regular target streams in most cases, when they were not informed about these stimuli *a priori*^13^. While the present experiment allows for a more precise, trial-by-trial evaluation of the perception, this comes at the expense of knowledge and task relevance of the target. It is therefore quite likely that the active representation of the target also played a role for its perception in the present study^48,49^. There is also no doubt that focused top-down attention enhances the detection of masked tones, as was demonstrated in the behavioral experiment in this study. Conversely, strong attentional distraction due to the presence of visual load may slightly reduce detection rate of those sounds, even without intramodal masking^50^. Therefore, and because the secondary-ear tones were task-relevant, we cannot rule out the possibility that some small amount of attention was devoted to them or that listeners didn’t switch their attention rapidly between ears during the task.

### Functional Dissociation of Left and Right AC Under Load

Hemispheric asymmetry is a central tenet of cognitive neuroscience. Traditionally, such asymmetry has been associated primarily with higher-order cognitive functions such as attention, where right-sided parietal lesions lead to hemispatial neglect^51,52^, and language, which is left dominant in approximately 95% and 70-85% of right- and left-handed individuals, respectively^53^. Though recent proposals have suggested more fundamental functional distinctions between the left and right hemispheres in both the visual^54^ and auditory^55–58^ domains, in audition, these were put forth primarily to account either for left lateralization of language and/or right lateralization of music. Furthermore, although many dichotic listening studies dating to the 1960s^59–61^ – including those conducted in split-brain patients^60,62,63^ – point to the importance of left-AC activity in auditory perceptual awareness, to our knowledge, the effects were only observed when using speech stimuli^59,64–66^.

Thus, our findings extend the notion of hemispheric asymmetry to potentially include basic perceptual awareness of nonverbal sound sequences, a finding that cannot be explained by current models. In particular, our findings suggest a strong association between auditory perceptual awareness and activity in left AC, at least under certain stimulus/task configurations. Support for this idea comes from recent studies employing similar multi-tone masker stimuli^18,67^ as well as a study examining the pre-stimulus network configuration of auditory cortex in a peri-threshold detection task^68^. That study found that for peri-threshold sounds (brief noise bursts) that were detected, connectivity indices between left and right AC increased and, critically, left AC acted as a network hub that was connected to distal brain areas in frontal and parietal cortex. However, their post-stimulus, noise-elicited responses did not show such asymmetry.

One interesting question going forward is whether the direction of the asymmetry we observed depends on hemispheric dominance. All but one of our participants were right-handed, and thus very likely left-dominant for language. The lone left-handed participant responded with her dominant hand, and her pattern of results were in line with the group average (i.e., left-AC predominance of awareness-related activity). However, as even most left-handed individuals are left-dominant for language^53^, she was likely also left-dominant, and testing the hypothesis that the direction of the asymmetry we observed depends on hemispheric dominance would require individualized language mapping.

Further questions posed by our findings are (i) the factors underlying the AAN asymmetry we observed, which are at present unclear but likely have to do with some aspect of the task utilized, and (ii) which aspects of functional brain organization enable such asymmetric processing of basic nonverbal sounds. Certain tasks have been shown to result in lateralized responses even when the same stimuli presented passively do not^69^. However, as for the split-brain research described above, this study used speech sounds. Studies using tonal stimuli have also observed asymmetries, which were attributed either to the mere presence of auditory distractors^70^ or by the task of detecting target sounds in the presence of distractors^71^.

The results from previous studies utilizing similar multi-tone cloud stimuli have been mixed, with some indicating little to no asymmetry^13,72,73^, and others showing left > right AAN responses^12,18,67^. Some of these studies used diotic and some dichotic presentation, suggesting that dichotic stimulation, per se, is not the cause of the asymmetry we observed. The only factor common to studies that observed asymmetry – and absent in those that didn’t – was the presence of task-relevant deviants in the target stream. Though this was not the case for one study^67^, there, the fact that only two tones comprised the target stream may have lead to them being processed as deviants. Interestingly, this was also the case for studies showing left > right asymmetries in other contexts^74,75^. Other recent work has found an effect of task difficulty on the lateralization of activity in auditory cortex^76^. Taken together, these studies suggest that the combination of auditory distraction and the difficulty of the segregation/discrimination task may be what underlies left lateralization across such diverse paradigms, supporting a role for left AC in “active listening”. This is supported by one recent study showing that lesions of left AC and adjacent peri-rolandic cortex were associated with worse performance on a rhythmic release from masking task^77^.

Yet, this does not explain the fact that in the present study, undetected targets, particularly those presented to the left ear, elicited robust negative responses, more strongly in right AC, in the same latency range as the N1/AAN. This stands in stark contrast to previous multi-tone masker studies, where virtually no activity was observed in the AAN latency range, in either hemisphere, in response to undetected targets, and if present, the N1 was never stronger than the P1^12^. One possible reason for the stronger N1 in the present case could be the sparser multi-tone masker compared to previous studies. Similar N1 responses as for undetected targets are observed for masker tones (Fig. S1), and the N1 was somewhat stronger for tones in the earlier part of the masker. In contrast to single-tone detection paradigms^15^, the difference in perceptual awareness is mainly the tone sequence in our case, whereas single tones may still be perceived but as part of the masker^78^. Accordingly, the left lateralization for the AAN observed here may be related to the perceptual organization of the target sequence in the context of a tone cloud with sparser masker tones.

However, while the masker-tone cloud was sparser for the secondary ear, the right-dominant N1 for undetected targets was even more consistent for primary-ear stimuli. One possibility could be an auditory-centric extension of right-lateralized attention networks^51,52^, and in particular the salience network, of which the right temporal parietal junction (TPJ) is a part. Irrespective of task-relevance, the TPJ typically responds to salient stimuli^51^. However, in the context of visual search tasks where suppression of task-irrelevant stimuli is important, TPJ activity in response to non-target items is suppressed^79,80^. Considering our paradigm in the context of search, if the right AC were coupled to the TPJ (not unlikely given their anatomical proximity and prior auditory findings^81^), we would expect right-AC activity to be suppressed during active search (for a target stream, in our case). This could explain why prior multitone masking studies have not observed such right-AC activity in response to undetected target streams^12,13^. However, in the context of the dichotic, dual-task used here, suppression of all non-relevant sound might impede detection of secondary-ear targets if attention is strongly focused towards the primary ear. Thus, having a more robust bottom-up representation of secondary-ear targets, which manifested here as a right-lateralized N1, potentially facilitates their perception. This hypothesis cannot be confirmed based on the data presented here due to the fact that several other factors differed between the present and previous paradigms, as discussed above.

## ACKNOWLEDGMENTS

This work was supported by the German Federal Ministry of Education and Research (01EV0712 to A.G.) and by Deutsche Forschungsgemeinschaft (DFG 593/5-1 to A.G.). The authors would like to thank Barbara Burghardt, Helmut Riedel, Esther Tauberschmidt, and Christian Uhlig for assistance with data collection as well as Andre Rupp and Helmut Riedel for helpful discussion.

## DATA AVAILABILITY STATEMENT

Stimuli, data, and software are available upon request.

## SUPPLEMENTAL FIGURES

**Figure S1.**
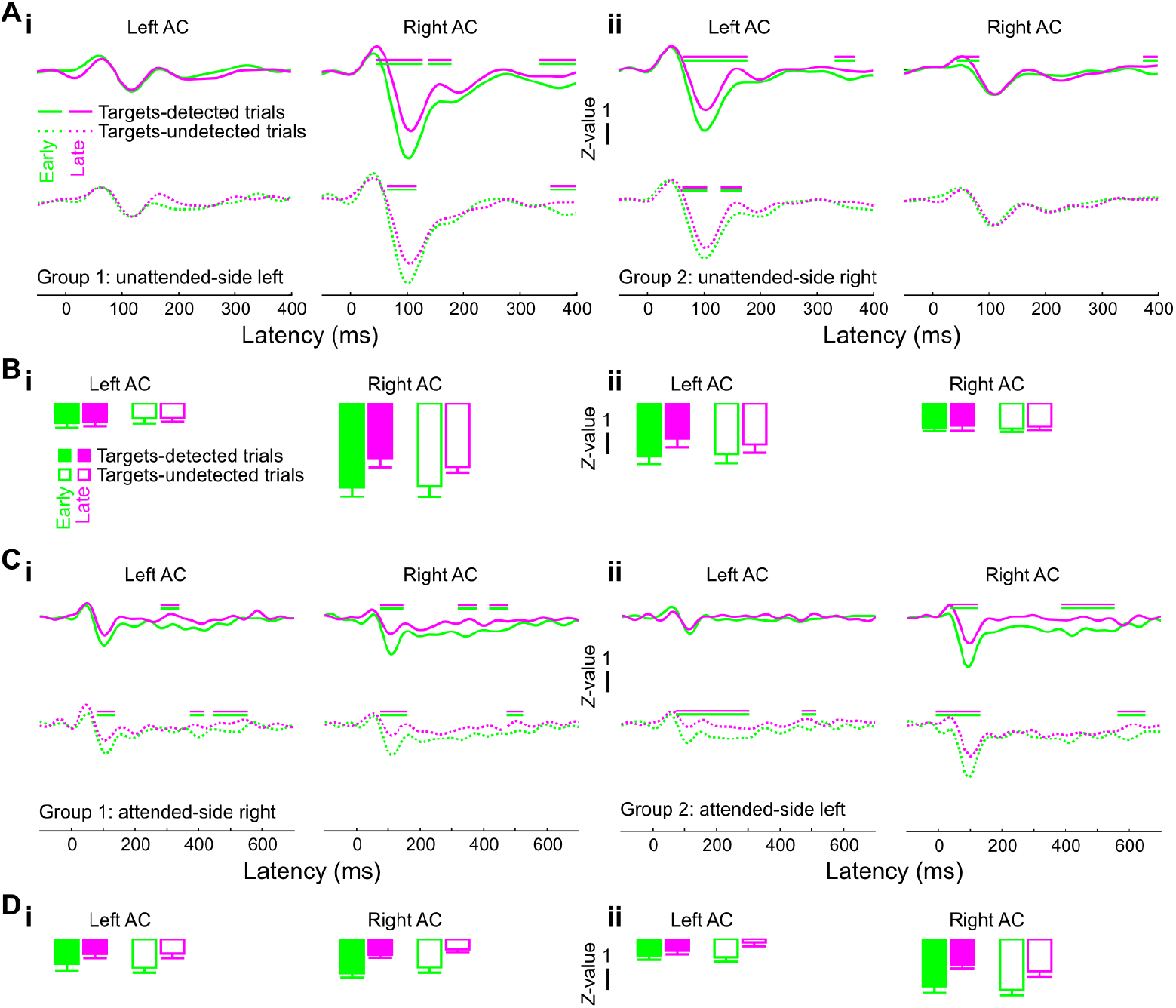
Masker-tone responses for primary- and secondary-ear maskers. **(A)** Grand-averaged timecourses for secondary-ear masker tones for each group separately for early (green) and late (magenta) masker events during targets-detected (solid traces) and targets-undetected (dotted) trials. Green/magenta bars above each panel denote statistically significant differences between the green and magenta traces. **(B)** Corresponding amplitudes during the N1 latency range. **(C)** Same as in (A), for primary-ear maskers. (D) Same as in (B), for primary-ear maskers.

## SUPPLEMENTAL TABLES

**Table S1.** Paired comparisons for secondary-ear hit rates sorted by target frequency and primary-ear percept.

|  | Group 1 |  |  | Group 2 |  |  |
| --- | --- | --- | --- | --- | --- | --- |
|  | 585 Hz | 914 Hz | 1430 Hz | 585 Hz | 914 Hz | 1430 Hz |
| Detected vs. Undetected | $T_{15} = -5.2, ***$ | $T_{15} = -1.0, n.s.$ | $T_{15} = -0.31, n.s.$ | $T_{15} = -1.0, n.s.$ | $T_{15} = -1.9, \cdot$ | $T_{15} = 0.60, n.s.$ |
| Undetected vs. Absent | $T_{15} = 4.5, ***$ | $T_{15} = -0.18, n.s.$ | $T_{15} = -0.25, n.s.$ | $T_{15} = -1.2, n.s.$ | $T_{15} = -0.65, n.s.$ | $T_{15} = -1.3, n.s.$ |
| Detected vs. Absent | $T_{15} = -1.3, n.s.$ | $T_{15} = -1.6, n.s.$ | $T_{15} = -1.0, n.s.$ | $T_{15} = -2.2, *$ | $T_{15} = -2.6, *$ | $T_{15} = -1.1, n.s.$ |

Significance codes: $\cdot = p < 0.1$ , $* = p < 0.05$ , $** = p < 0.01$ , $*** = p < 0.001$ .

**Table S2.** Primary-ear hit- and false-alarm rates, d’, and statistics, sorted by secondary-ear percept and tone position.

|  |  | Tone position |  |  |  |  |  |  |  |  |
| --- | --- | --- | --- | --- | --- | --- | --- | --- | --- | --- |
|  |  | 1 | 2 | 3 | 4 | 5 | 6 | 7 | 8 | 9 |
| TPR | Absent | $0.08 \pm 0.02$ | $0.32 \pm 0.03$ | $0.49 \pm 0.03$ | $0.60 \pm 0.03$ | $0.69 \pm 0.02$ | $0.73 \pm 0.02$ | $0.77 \pm 0.02$ | $0.78 \pm 0.02$ | $0.78 \pm 0.02$ |
| | Undetected | $0.08 \pm 0.02$ | $0.31 \pm 0.03$ | $0.49 \pm 0.02$ | $0.61 \pm 0.02$ | $0.69 \pm 0.02$ | $0.75 \pm 0.02$ | $0.78 \pm 0.02$ | $0.79 \pm 0.02$ | $0.79 \pm 0.02$ |
| | Detected | $0.11 \pm 0.02$ | $0.38 \pm 0.03$ | $0.52 \pm 0.03$ | $0.60 \pm 0.03$ | $0.64 \pm 0.03$ | $0.67 \pm 0.03$ | $0.69 \pm 0.03$ | $0.70 \pm 0.03$ | $0.70 \pm 0.03$ |
| | $F_{2,30}, p$ | $0.9, n.s.$ | $1.8, n.s.$ | $0.4, n.s.$ | $0.0, n.s.$ | $1.1, n.s.$ | $3.0, \cdot$ | $4.3, *$ | $4.4, *$ | $4.4, *$ |
| FPR | Absent | $0.01 \pm 0.00$ | $0.02 \pm 0.01$ | $0.03 \pm 0.01$ | $0.04 \pm 0.01$ | $0.05 \pm 0.01$ | $0.06 \pm 0.01$ | $0.07 \pm 0.01$ | $0.08 \pm 0.01$ | $0.08 \pm 0.01$ |
| | Undetected | $0.00 \pm 0.00$ | $0.02 \pm 0.01$ | $0.03 \pm 0.01$ | $0.04 \pm 0.01$ | $0.06 \pm 0.01$ | $0.08 \pm 0.01$ | $0.10 \pm 0.02$ | $0.10 \pm 0.02$ | $0.10 \pm 0.02$ |
| | Detected | $0.01 \pm 0.00$ | $0.02 \pm 0.01$ | $0.03 \pm 0.01$ | $0.04 \pm 0.01$ | $0.04 \pm 0.01$ | $0.05 \pm 0.01$ | $0.06 \pm 0.01$ | $0.06 \pm 0.01$ | $0.06 \pm 0.01$ |
| | $F_{2,30}, p$ | $0.6, n.s.$ | $0.2, n.s.$ | $0.0, n.s.$ | $0.2, n.s.$ | $1.0, n.s.$ | $1.3, n.s.$ | $2.0, n.s.$ | $2.1, n.s.$ | $2.1, n.s.$ |
| $d'$ | Absent | $0.6 \pm 0.12$ | $1.6 \pm 0.12$ | $1.9 \pm 0.09$ | $2.1 \pm 0.10$ | $2.3 \pm 0.10$ | $2.3 \pm 0.11$ | $2.4 \pm 0.10$ | $2.3 \pm 0.10$ | $2.3 \pm 0.10$ |
| | Undetected | $0.7 \pm 0.11$ | $1.6 \pm 0.09$ | $2.0 \pm 0.09$ | $2.1 \pm 0.10$ | $2.2 \pm 0.10$ | $2.2 \pm 0.10$ | $2.2 \pm 0.11$ | $2.3 \pm 0.11$ | $2.3 \pm 0.11$ |
| | Detected | $0.8 \pm 0.10$ | $1.7 \pm 0.10$ | $2.0 \pm 0.11$ | $2.1 \pm 0.12$ | $2.2 \pm 0.12$ | $2.3 \pm 0.11$ | $2.2 \pm 0.11$ | $2.3 \pm 0.11$ | $2.3 \pm 0.11$ |
| | $F_{2,30}, p$ | $0.5, n.s.$ | $0.4, n.s.$ | $0.0, n.s.$ | $0.0, n.s.$ | $0.5, n.s.$ | $0.2, n.s.$ | $0.6, n.s.$ | $0.1, n.s.$ | $0.1, n.s.$ |

TPR = True-positive rate. FPR = False-positive rate. Significance codes: $\cdot = p < 0.1$ , $* = p < 0.05$ , $** = p < 0.01$ , $*** = p < 0.001$ .

**Table S3.** Secondary-ear peak-value statistics for the left hemisphere as a function of group and target-tone position.

|  |  | Group 1 |  | Group 2 |  | Groups 1 & 2 |  |
| --- | --- | --- | --- | --- | --- | --- | --- |
|  |  | 80-160 ms | 160-240 ms | 80-160 ms | 160-240 ms | 80-160 ms | 160-240 ms |
| Left AC | Position 1 | $T_{15} = -1.3, n.s.$ | $T_{15} = -0.53, n.s.$ | $T_{15} = -1.9, \cdot$ | $T_{15} = -0.39, n.s.$ | $T_{31} = -2.3, *$ | $T_{31} = -0.66, n.s.$ |
| | Position 2 | $T_{15} = -1.0, n.s.$ | $T_{15} = -1.2, n.s.$ | $T_{15} = -1.6, n.s.$ | $T_{15} = -3.3, **$ | $T_{31} = -1.8, \cdot$ | $T_{31} = -3.1, **$ |
| | Position 3 | $T_{15} = -1.9, \cdot$ | $T_{15} = -2.9, *$ | $T_{15} = -4.2, ***$ | $T_{15} = -3.9, **$ | $T_{31} = -4.2, ***$ | $T_{31} = -4.9, ***$ |
| | Position 4 | $T_{15} = -3.0, **$ | $T_{15} = -2.1, \cdot$ | $T_{15} = -5.3, ***$ | $T_{15} = -3.9, **$ | $T_{31} = -5.6, ***$ | $T_{31} = -4.1, ***$ |
| Right AC | Position 1 | $T_{15} = -0.40, n.s.$ | $T_{15} = -0.82, n.s.$ | $T_{15} = -0.10, n.s.$ | $T_{15} = 0.74, n.s.$ | $T_{31} = -0.35, n.s.$ | $T_{31} = -0.25, n.s.$ |
| | Position 2 | $T_{15} = -0.23, n.s.$ | $T_{15} = 0.24, n.s.$ | $T_{15} = -1.0, n.s.$ | $T_{15} = 0.10, n.s.$ | $T_{31} = -0.77, n.s.$ | $T_{31} = 0.26, n.s.$ |
| | Position 3 | $T_{15} = -1.7, n.s.$ | $T_{15} = 0.05, n.s.$ | $T_{15} = 0.12, n.s.$ | $T_{15} = -0.39, n.s.$ | $T_{31} = -1.2, n.s.$ | $T_{31} = -0.29, n.s.$ |
| | Position 4 | $T_{15} = -1.2, n.s.$ | $T_{15} = -0.17, n.s.$ | $T_{15} = -0.50, n.s.$ | $T_{15} = 0.12, n.s.$ | $T_{31} = -1.2, n.s.$ | $T_{31} = -0.06, n.s.$ |

Significance codes: $\cdot = p < 0.1$ , $* = p < 0.05$ , $** = p < 0.01$ , $*** = p < 0.001$ . Right hemisphere: group 1 ( $T_{15} \leq 1.7, p \geq 0.12$ ), group 2 ( $T_{15} \leq 1.0, p \geq 0.32$ ), groups 1 & 2 ( $T_{31} \leq 1.2, p \geq 0.24$ ).

**Table S4.** Primary-ear peak-value statistics sorted by group, time window, and hemisphere.

|  | Group 1 |  | Group 2 |  | Groups 1 & 2 |  |
| --- | --- | --- | --- | --- | --- | --- |
|  | 80-160 ms | 160-240 ms | 80-160 ms | 160-240 ms | 80-160 ms | 160-240 ms |
| Left AC | $T_{15} = -4.3, ***$ | $T_{15} = -5.6, ***$ | $T_{15} = -3.8, **$ | $T_{15} = -4.6, ***$ | $T_{31} = -5.5, ***$ | $T_{31} = -7.1, ***$ |
| Right AC | $T_{15} = -0.30, n.s.$ | $T_{15} = -1.6, n.s.$ | $T_{15} = -0.79, n.s.$ | $T_{15} = -2.8, *$ | $T_{31} = -0.79, n.s.$ | $T_{31} = -3.0, **$ |

Significance codes: $\cdot = p < 0.1$ , $* = p < 0.05$ , $** = p < 0.01$ , $*** = p < 0.001$ .

