## supplemental audio files for "Hemispheric decoupling of awareness-related activity in human auditory cortex under informational masking and divided attention"

**Audio S1. Example of an attended-side stimulus sequence.** A regularly repeating target-tone stream at 1430 Hz with an SOA of 800 ms is embedded within a random multi-tone masker "cloud". The target stream can initially be quite difficult to identify (headphones are strongly recommended), but is typically heard much more readily after listening to a filtered version in which the loudness of the masker tones is suppressed (Audio S2). Also note that two of the target tones near the end of the sequence were amplitude-modulated, which listeners were instructed to count if and when they were aware of the target stream.

**Audio S2. Filtered version of Audio S1.** Same as Audio S1, but with a band-pass filter applied to a narrow range around the frequency (1430 Hz) of the target. Here, the isochronous target stream is much easier to identify.

**Audio S3. Example of an unattended-side stimulus sequence.** Four identical target tones with a frequency of 585 Hz and SOA of 400 ms are placed near the middle of the sequence. Here, the masker cloud used is much sparser than for the attended side, making the targets easier to identify without the presence of the attended-side sequence. However, if the targets are not readily identified, see Audio S4.

**Audio S4. Filtered version of Audio S3.** Same as Audio S3, but with a band-pass filter applied to a narrow range around the frequency (585 Hz) of the target, making the isochronous target stream much easier to identify.

**Audio S5. Example of an actual stimulus used in the study.** Both attended- and unattended-side sequences are present (Audio S1 and Audio S3, respectively). Here, the right (left) ear was designated as the attended (unattended), as in Group 1, though this was flipped in Group 2. Also, note that all combinations of attended- and unattended-side target-stream presence and target-stream frequencies were equally likely so as to keep each side uninformative about the other.
